# Marine Dadabacteria exhibit genome streamlining and phototrophy-driven niche partitioning

**DOI:** 10.1101/2020.06.22.165886

**Authors:** Elaina D. Graham, Benjamin J. Tully

**Author notes:** authors contributed equally.

## Abstract

The remineralization of organic material via heterotrophy in the marine environment is performed by a diverse and varied group of microorganisms that can specialize in the type of organic material degraded and the niche they occupy. The marine *Dadabacteria* are cosmopolitan in the marine environment and belong to a candidate phylum for which there has not been a comprehensive assessment of the available genomic data to date. Here in, we assess the functional potential of the oligotrophic, marine *Dadabacteria* in comparison to terrestrial, coastal, and subsurface members of the phylum. Our analysis reveals that the marine *Dadabacteria* have undergone a genome streamlining event, reducing their genome size and the nitrogen content of their DNA and predicted proteome, relative to their terrestrial counterparts. Collectively, the *Dadabacteria* have the potential to degrade microbial particulate organic matter, specifically peptidoglycan and phospholipids. The marine *Dadabacteria* belong to two clades with distinct ecological niches in global metagenomic data: a shallow clade with the potential for photoheterotrophy through the use of proteorhodopsin, present predominantly in surface waters up to 100m depth; and a deep clade lacking the potential for photoheterotrophy that is more abundant in the deep photic zone.

## Introduction

Heterotrophy in the marine environment is a complex process with many organisms contributing to the remineralization of organic matter. In the surface ocean, ∼50% of new organic carbon is remineralized by heterotrophs within the first 100 meters (Cole *et al*., 1988; Ducklow *et al*., 1993). Despite the importance of this process to the overall ocean carbon budget, the specific contributions of the phylogenetically diverse marine bacterioplankton community remain poorly constrained. The metabolic capacity of the community members directly governs the types of organic material that can be degraded in a particular environment (Steen *et al*., 2019). Heterotrophs occupy a spectrum of metabolic diversity and growth strategies (Malik *et al*., 2019). While copiotrophs exploit multiple organic resources and/or undergo rapid growth in response to nutrient availability, oligotrophs specialize in a limited number of resources and dominate in low nutrient environments (Vergin *et al*., 2013). Because of the interplay of heterotrophs on this spectrum of metabolic diversity, it is important to understand the role(s) that specific groups play in the degradation of organic matter in the surface ocean.

An evolutionary feature that is common among marine oligotrophs is the reduction and simplification of the genome. This evolutionary trajectory has been posited as the theory of genome streamlining, in which organisms that grow in nutrient limited environments undergo selection to reduce cellular demand for specific compounds and nutrients (Giovannoni *et al*., 2014). While originating in the marine environment (Rocap *et al*., 2003; Giovannoni *et al*., 2005), genome streamlining has been identified in numerous habitats for a variety of microorganisms (Luo *et al*., 2014; Castelle *et al*., 2015; Brewer *et al*., 2016; Neuenschwander *et al*., 2018). While reduced genome size can be a result of genome streamlining, several other genome modifications can interact to reduce the overall cellular demand for nutrients, including an increase in coding density and the absence of paralogs/gene duplication events, reducing the total amount of DNA that must undergo replication during cell division, and, in nitrogen limited environments, a reduction in contribution of nitrogen to both the DNA and the proteome (Getz *et al*., 2018). The theory of genome streamlining is an important avenue for understanding microbiology and provides important insights into the evolutionary history and ecological distributions of a microorganism.

Here in, we assess the potential contributions of the *Dadabacteria* to marine heterotrophy. A phylum level group, the *Dadabacteria* (formerly SBR1093) lack a cultured representative and have not been extensively assessed for their potential contributions to biogeochemical cycles though they have been detected in numerous terrestrial and marine environments. The first *Dadabacteria* genome was reconstructed from industrial active sludge and reported to possess the capacity for carbon fixation through the 3-hydroxybutyrate/4-hydroxypropionate cycle (Wang *et al*., 2014). Interestingly, multiple *Dadabacteria* metagenome-assembled genomes were reconstructed from the *Tara* Oceans global, marine metagenomic samples, though their exact role in the marine environment was unknown (Tully *et al*., 2018; Parks *et al*., 2017; Delmont *et al*., 2018). Our analysis reveals that the marine *Dadabacteria* are likely heterotrophic oligotrophs that have undergone genome streamlining with the capacity to degrade microbially derived peptidoglycan as a carbon source with further metabolic diversification between shallow and deep-water niches.

## Materials and methods

### Collect, assess and clean genomes, and construct phylogenomic trees

MAGs generated from several studies using the *Tara* Oceans metagenomics dataset were initially identified as *Dadabacteria* based on 16S rRNA phylogeny and 16 concatenated ribosomal proteins (ribosomal proteins L2, L3, L4, L5, L6, L14, L16, L18, L22, L24, S3, S8, S10, S17, and S19) (Hug *et al*., 2016). All *Dadabacteria* genomes identified in NCBI (as of August 2019) (Anantharaman *et al*., 2016; Hug *et al*., 2019; Kato *et al*., 2018; Zhou *et al*., 2020; Ward *et al*., 2019) and one *Dadabacteria* genome (formally Candidate Phylum SBR1093) derived from Wang *et al*. (2014) was also included. Genomes reconstructed from Tully *et al*. (2018) were subjected to manual assessments for quality using the same methodology as in Graham *et al*. (Graham *et al*., 2018). Briefly, read coverage and DNA compositional data was utilized to bin additional contigs (>5kb) from the *Tara* Oceans Longhurst province the original *Dadabacteria* MAG was reconstructed from using CONCOCT (v.0.4.1; parameters: -c 800 -I 500) (Alneberg *et al*., 2014). To improve completion estimates overlapping CONCOCT and BinSanity bins were visualized in Anvi’o (Eren *et al*., 2015) and manually refined to minimize contamination estimates and improve genome completion. Genomes from Delmont *et al*. (2018) were also visualized in Anvi’o and manually curated based on DNA composition (%G+C and tetranucleotide frequencies) to minimize contamination estimates and improve genome completion.

### Dadabacteria

MAGs were assessed for quality through the PhyloSanity workflow of the tool MetaSanity. Estimated completeness, contamination, and strain heterogeneity were determined using CheckM (v1.0.18) (Parks *et al*., 2015). The estimated completeness and MAG size were used to calculate an approximate genome size for the complete genome. Additionally, the CheckM QA workflow was used to calculate the coding density. Phylogeny was confirmed using GTDB-Tk (v1.0.0; database ver. 89; parameters: classify_wf defaults) (Chaumeil *et al*., 2019). The GTDB-Tk de novo workflow was used to construct a multiple sequence alignment (MSA) of the *Dadabacteria* MAGs using the bac120 marker set and with f_SZUA-79 set as the outgroup. The full MSA was reduced to include the following lineages related to the *Dadabacteria*: SZUA-79, *Chrysiogenetota, Deferribacterota, Thermosulfidibacterota, Aquificota, Camplyobacterota*. The MSA was refined using MUSCLE (v3.8.31, parameter: - refine) (Edgar, 2004) and FastTree (v2.1.10, parameters: -lg, -gamma) (Price *et al*., 2010) was used to generate a phylogenetic tree that was visualized using the Interactive Tree of Life (IToL) (Letunic and Bork, 2016).

### Functional annotation

For functional annotation and evidence of genomic streamlining, due to the limited number of available MAGs, all genomes were considered during the analysis. *Dadabacteria* MAGs were assessed for putative metabolic functionality through the FuncSanity workflow of the tool MetaSanity (Neely *et al*., 2019). All downstream analysis uses the of putative CDS as predicted by Prokka (v1.13.3) (Seemann, 2014). Putative CDS were assigned to carbohydrate-active enzyme (CAZy) families based on HMM models from dbCAN (v6) (Yin *et al*., 2012) using hmmsearch (v3.1b2; parameter: -T 75) (Finn *et al*., 2011). The output from MetaSanity that combines the CAZy matches for all submitted genomes (MetaSanity output file: combined.cazy) was used to determine the number of CAZy matches per Mbp in each MAG, including a curated selection of glycoside hydrolases (GH) and carbohydrate-binding module (CBM) containing proteins, excluding matches to CAZy subfamily HMM models (e.g., matches to GH13 model were included, while matches to GH13_9 model, etc. were excluded).

CDS were determined to be putative peptidases through hmmsearch (parameter: -T 75) using PFAM (El-Gebali *et al*., 2018) HMM models selected to represent the MEROPS families (Rawlings *et al*., 2013). Putative peptidases were screened for signatures denoting possible extracellular localization using PSORTb (v3.0) (Yu *et al*., 2010) and SignalP (v4.1) (Petersen *et al*., 2011). First, PSORTb was used to identify all putative peptidases with the localization assignment of “extracellular”, “cellwall”, or “unknown”. For any putative peptidase that had “unknown” localization, if SignalP predicted a transmembrane helix, the peptidase was determined to be putatively extracellular.

Metabolic functions of interest were identified based on the KEGG-Decoder (Graham *et al*., 2018) output (v1.0.10) as implemented in MetaSanity (MetaSanity output file: KEGG.final.tsv). As part of this workflow, CDS were assigned to KEGG Ontology (KO) identifiers using KofamScan (v1.2.0) (Aramaki *et al*., 2020) and the accompanying KOfam HMM models and then assigned to a set of manually curated pathways and processes. Additionally, metabolisms of interest, especially those lacking KOfam HMM models, were searched independently and incorporated using KEGG-Expander as implemented in MetaSanity.

Additional databases were used to identify feature of interests within the *Dadabacteria* MAGs. Putative metabolic functions of interest shared between the four phylogenetic clades were identified using eggNOG-mapper (Huerta-Cepas *et al*., 2017) (http://eggnog-mapper.embl.de/; default parameters for “Auto adjust per query”) and precomputed eggNOG clusters (v5.0) (Huerta-Cepas *et al*., 2018). antiSMASH (v5.0.0) (Blin *et al*., 2019) was used to detect secondary metabolite biosynthetic gene clusters (parameters: --cb-general --cb-knownclusters --cb-subclusters --asf --pfam2go --smcog-trees). Based on matches to the rhodopsin PFAM HMM model (PF01036) performed as part of the KEGG-Decoder analysis, putative rhodopsin CDS were compared to the MicRhoDE database (Boeuf *et al*., 2015) using BLASTP (Camacho *et al*., 2009) (http://application.sb-roscoff.fr/micrhode/doblast; default parameters for “All Micrhode” option) and assigned to a previously identified classes based on the highest scoring result. Additionally, putative rhodopsins were aligned with MUSCLE (parameter: -iter 8) and the 17 amino acid (aa) region that contains the crucial aa for determining function (aa site 97 & 108) and spectral tuning (aa site 105) were categorized based on known rhodopsin relationships.

### Genomic streamlining

Putative CDS were used to calculate the total number of carbon and nitrogen atoms present in the predicted proteome and the corresponding ratio of each MAG (https://github.com/edgraham/CNratio). For identifying duplicate genes in a MAG, first, all putative CDS in a MAG was compared against each other using DIAMOND BLASTP (Buchfink *et al*., 2014) (parameters: --more-sensitive –max-taget-seqs 300). BLAST matches were filtered using the minbit approach described in (Benedict *et al*., 2014), where significant matches were determined based on the relative comparison of bitscore values. Minbit was calculated for protein A compared to protein B as

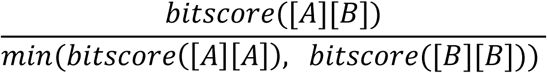

retaining all BLAST matches ≥0.5. BLAST matches above this threshold were reformatted and clustered using MCL (van Dongen and Abreu-Goodger, 2011) (mcxload parameters: --abc -- stream-mirror --stream-neg-log10 -stream-tf ceil(200); mcl default parameters; mcxdump parameter: -icl). All clusters in the mcxdump output were considered to be gene duplication events within the MAG.

### Ecological distribution and environmental correlations

For determining the ecological distribution and environmental correlations, a non-redundant set of MAGs was determined using FastANI (Jain *et al*., 2018) (v1.3; parameters: --frag-length 1500) with a representative selected from a cluster of genomes with ≥98.5% average nucleotide identity. Metagenomes derived from bioGEOTRACES (Biller *et al*., 2018) (bGT) and *Tara* Oceans (Sunagawa *et al*., 2015) were mapped against the non-redundant set of *Dadabacteria* genomes using bowtie2 (Langmead and Salzberg, 2012) (v2.3.4.1, parameters: -q, --no-unal), converted from a SAM to BAM file using samtools (Li *et al*., 2009) (v.1.9; view; sort), and filtered using BamM (v1.7.0, parameters: -- percentage_id 0.95, --percentage_aln 0.75). featureCounts (Liao *et al*., 2014) (v1.5.3, default parameters) implemented through Binsanity-profile (Graham *et al*., 2017) (v0.3.3, default parameters) was used to generate read counts for each contig from the filtered BAM files. Read counts were used to calculate the relative fraction of each genome in the sample (Eq. 1) and determine the reads per kbp of each genome per Mbp of metagenomic sample (RPKM) (Eq. 2).

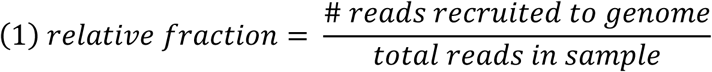

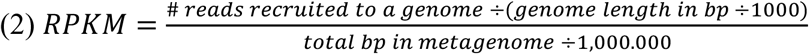

Environmental data was accessed from GEOTRACES Intermediate Data Product 2017 (Version 2) (Schlitzer *et al*., 2018) and paired with the corresponding metagenome sample ID. In many cases there were multiple CTD casts associated with a particular station and depth. The mean value was used in cases where a parameter was measured multiple times at the same depth and station. Environmental data was paired with a metagenome only if the depth was within one meter of the metagenome. RPKM values for *Dadabacteria* genomes from all samples with available environmental data were used in a canonical correspondence analysis (CCA) in Past4 (Hammer *et al*., 2001) (v.4.01; trioplot amp 1.5, scaling type 2). Only environmental data that was measured for ≥90% of the samples were used to perform the CCA. RPKM values were normalized (log(n+1)) prior to CCA. Transect plots were made in Ocean Data View (v5.2.1; DIVA Gridding; *Schlitzer, Reiner*, Ocean Data View, https://odv.awi.de, 2020). Bathymetry was pulled from General Bathymetric Chart of the Oceans (GEBCO 2014; doi: 10.1594/PANGAEA.708081).

## Results and discussion

As a candidate phylum, understanding the ecological role of the *Dadabacteria* remains difficult as only 48 MAGs are currently available for analysis (available August 2019). Based on the phylogenetic reconstruction (Figure 1A; Supplemental Table 1), the phylum partitions into three distinct clades from three predominant environmental sources: terrestrial hot springs, “terrestrial” (which includes MAGs from the terrestrial subsurface, oil-polluted marine systems, marine sponges, marine sediment, and hydrothermal vents), and the marine environment. Within the “marine clade”, there are two distinct subclades. Evidence from the predicted metabolisms and the ecological distributions of the marine clade support the demarcation of a surface and deep phylogenetic clade (see below). The marine clades harbor genomic features that differentiate them from the predominantly terrestrial clades, specifically with regards to genomic evolutionary selection (*e*.*g*., streamlining) and putative metabolisms.

**Figure 1.**
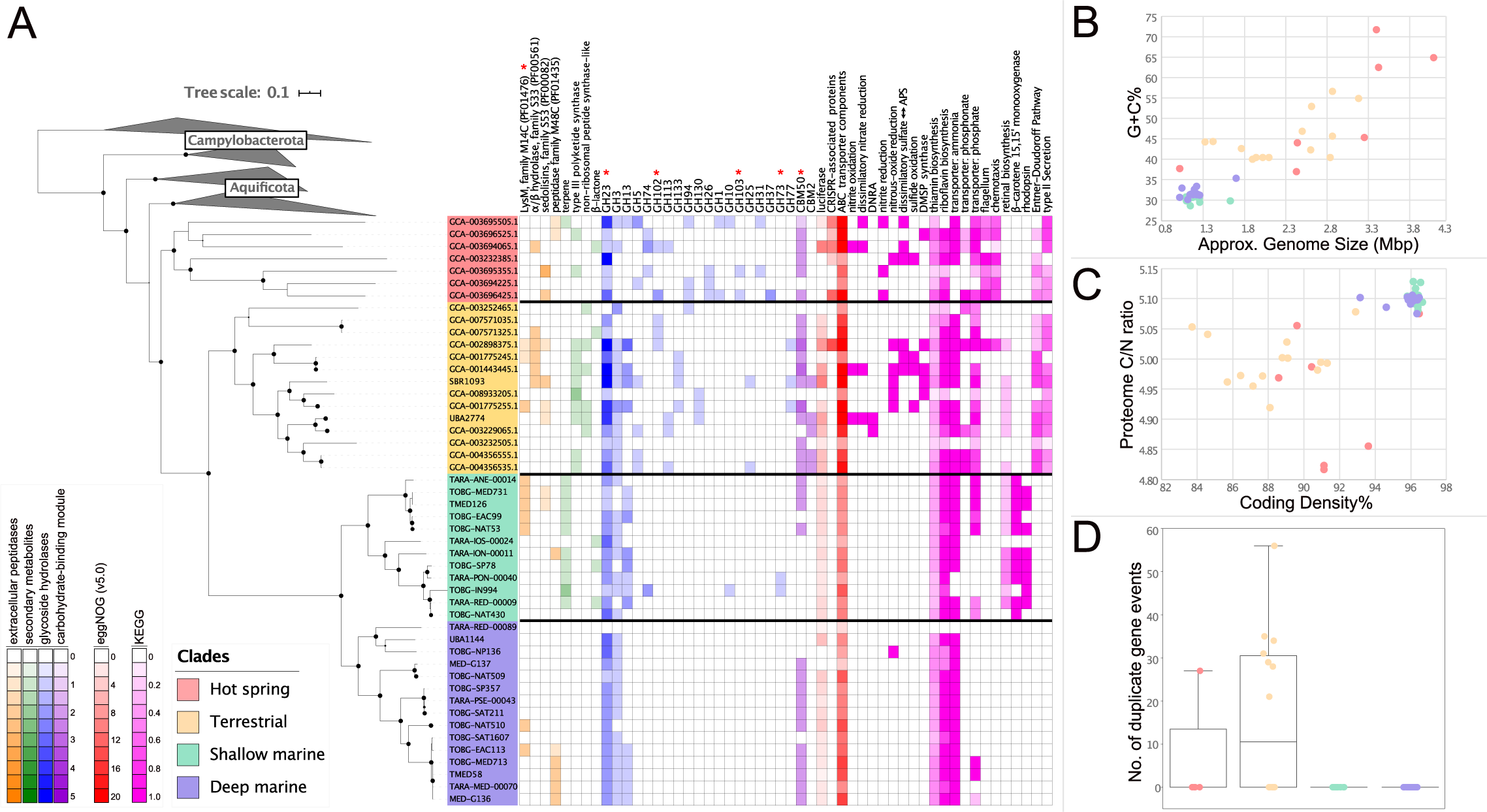
A. A phylogenomic tree of the bac120 marker set for the *Dadabacteria* and related phyla and a heatmap displaying functions of interest for each *Dadabacteria* MAG. Putative extracellular peptidase, secondary metabolite, glycoside hydrolase, and carbohydrate-binding module counts are displayed on a scale from 0-5. Functions inferred from eggNOG counts are displayed on a scale from 0-20+. Metabolic processes inferred from KEGG are displayed on a scale for 0-1, as a fraction of a particular metabolism detected. B. A scatter plot of percent G+C (%G+C) and approximate complete genome size in megabase pairs (Mbp) for each *Dadabacteria* MAG. C. A scatterplot of putative proteome carbon-to-nitrogen content ratio and percent coding density for each *Dadabacteria* MAG. D. The number of duplicate gene events in each *Dadabacteria* MAG.

The marine *Dadabacteria* have undergone a genome streamlining process in comparison to the terrestrial and hot spring lineages. The marine *Dadabacteria* exhibit all five traits associated with genome streamlining: reduced genome size, decreased %GC content, increased C/N ratio in the predicted proteome, increased coding density, and limited/no gene duplication events (Figure 1B-D; Supplemental Table 1) (Getz *et al*., 2018; Giovannoni *et al*., 2014). The estimated complete marine *Dadabacteria* genome is ∼1.22Mb with >96% coding density, smaller in size and similar in coding density to the well-studied oligotrophic SAR11 clade (Giovannoni *et al*., 2005; Grote *et al*., 2012). The presence of the *Dadabacteria* MAGs reconstructed from multiple oligotrophic *Tara* Oceans regions would suggest that these organisms, like other oligotrophs, are adapted to environments with low nutrient concentrations (Supplemental Figure 1). Modifications in GC content and proteome C/N ratio are associated with lowering the nitrogen demand for organisms in nitrogen-limited environments. While small genomes, devoid of paralogs and with high coding density, are thought to have reduced energy requirements for division and growth. These genomic modifications may provide the *Dadabacteria* with an advantage in oligotrophic marine environments and provide further evidence that the theory of genome streamlining is a common evolutionary response to nutrient limitation in the environment.

While the SBR1093 MAG was implicated in carbon fixation via the 3-hydroxypriopinate/4-hydroxybutyrate cycle (Wang *et al*., 2014), analysis of the *Dadabacteria* phylum reveals, especially for the marine clades, a predominantly heterotrophic lifestyle (Figure 1A). Except for the SBR1093 MAG, no other *Dadabacteria* MAGs have the potential for carbon fixation (Supplemental Table 2). Several MAGs from the hot spring and terrestrial clades have the potential to interface with the nitrogen and sulfur cycles with metabolic processes involved in denitrification, dissimilatory nitrate reduction to ammonia (DNRA), sulfate reduction, sulfide oxidation, and the production of dimethylsulfoniopropionate (DMSP) (Figure 1A). However, the marine clades lack these particular metabolic pathways, while maintaining a heterotrophic potential for proteins and complex carbohydrates, including starch/glycogen (β-glucosidase and α-amylase). One consistent target for the extracellular peptidases (LysM) and carbohydrate-active enzymes (CAZymes; peptidoglycan lyase and CBM Family 50) across the *Dadabacteria* clades is peptidoglycan, the polymer of the microbial cell wall. It may be possible that these predicted proteins are responsible for the internal recycling of the cell wall during cell division or an indication that the *Dadabacteria* occupy a niche capable of recycling microbially derived particulate organic matter (POM).

Interestingly, the number extracellular peptidases, CAZymes, and ATP-binding cassette-type (ABC-type) transporter components normalized for MAG length across all four clades remains consistent even as the overall diversity within each group of proteins decreases (Figure 1A; Supplemental Tables 3-5). This may highlight an interplay between heterotrophic metabolic diversity and substrate utilization efficiency as *Dadabacteria* genome size decreases during streamlining. Additionally, there are several other metabolic processes that distinguish the four clades and highlight the divide between the terrestrial clades and marine clades. Specifically, for the hot spring clade, the prevalence of CRISPR-associated proteins (used as proxy for CRISPR arrays due low recovery in MAGs), motility, and chemotaxis suggesting that both avoidance of viral predation and physical adjustments within the hot spring environment are important evolutionary advantages (Supplemental Tables 2 & 5). Distinct for the two terrestrial clades, are the presence of phosphonate and phosphate ABC transporters, the Entner-Doudoroff pathway, an alternative pathway to glycolysis for glucose degradation, and a Type II secretion system (Supplemental Tables 2 & 6). In many marine systems, phosphorous, like nitrogen, can be a limiting resource. All four clades possess ABC-type phospholipid transporters (Supplemental Table 6), so while most of the marine clades lack phosphonate and phosphate transporters, the presence of phospholipid transporters suggest these organisms may recover phosphorous for cellular demand from POM.

The shallow and deep marine clades have several distinguishing metabolic properties. Potentially most importantly are the mechanisms related to utilizing light energy. Uniquely amongst the *Dadabacteria*, the surface clade possesses rhodopsins and the biosynthetic capacity for retinal synthesis (Figure 1A). Based on the present amino acids, it is predicted that all of the identified rhodopsins are H^+^-pumping proteorhodopsins (Supplemental Table 7). For the eight identified proteorhodopsins within this clade, all but one are predicted to be spectrally tuned to absorb blue light (Supplemental Table 7). The surface clade also has the capacity to produce terpene secondary metabolites (Supplemental Table 8). Terpenes are organic hydrocarbons that have been shown to be associated with carotenoid synthesis (Gershenzon and Dudareva, 2007). These terpenes may be related to the production of β-carotene, a biological precursor to retinal, or to production of other unidentified carotenoids (Supplemental Table 6). The deep clade lack proteorhodopsins, retinal biosynthesis, and terpene secondary metabolites (Figure 1A). Within the deep clade, the presence of starch/glycogen and peptidoglycan degradation mechanisms may suggest that these heterotrophic processes are the predominant avenues for energy.

The metabolic division based on the utilization of light via proteorhodopsins between the shallow and deep clades is reflected in the ecological distribution of a clades. Using a nonredundant set of the marine *Dadabacteria* MAGs, the large global metagenomic datasets (*Tara* Oceans and bGT) were mapped against the MAGs and used to assess where the *Dadabacteria* occurred through the water column (Supplemental Tables 9 & 10). The two datasets have distinct properties that allow for varying perspectives on the ecology of the *Dadabacteria. Tara* Oceans is globally distributed with multiple size fractions and depths into the mesopelagic, while bGT provides several high-resolution cruise tracks with multiple depths between the surface and ∼250m depth. The results from *Tara* Oceans demonstrate that, broadly, the marine clades are present in the planktonic size fraction (<3μm) and almost exclusively found in the epipelagic (Supplemental Figure 2).

**Figure 2.**
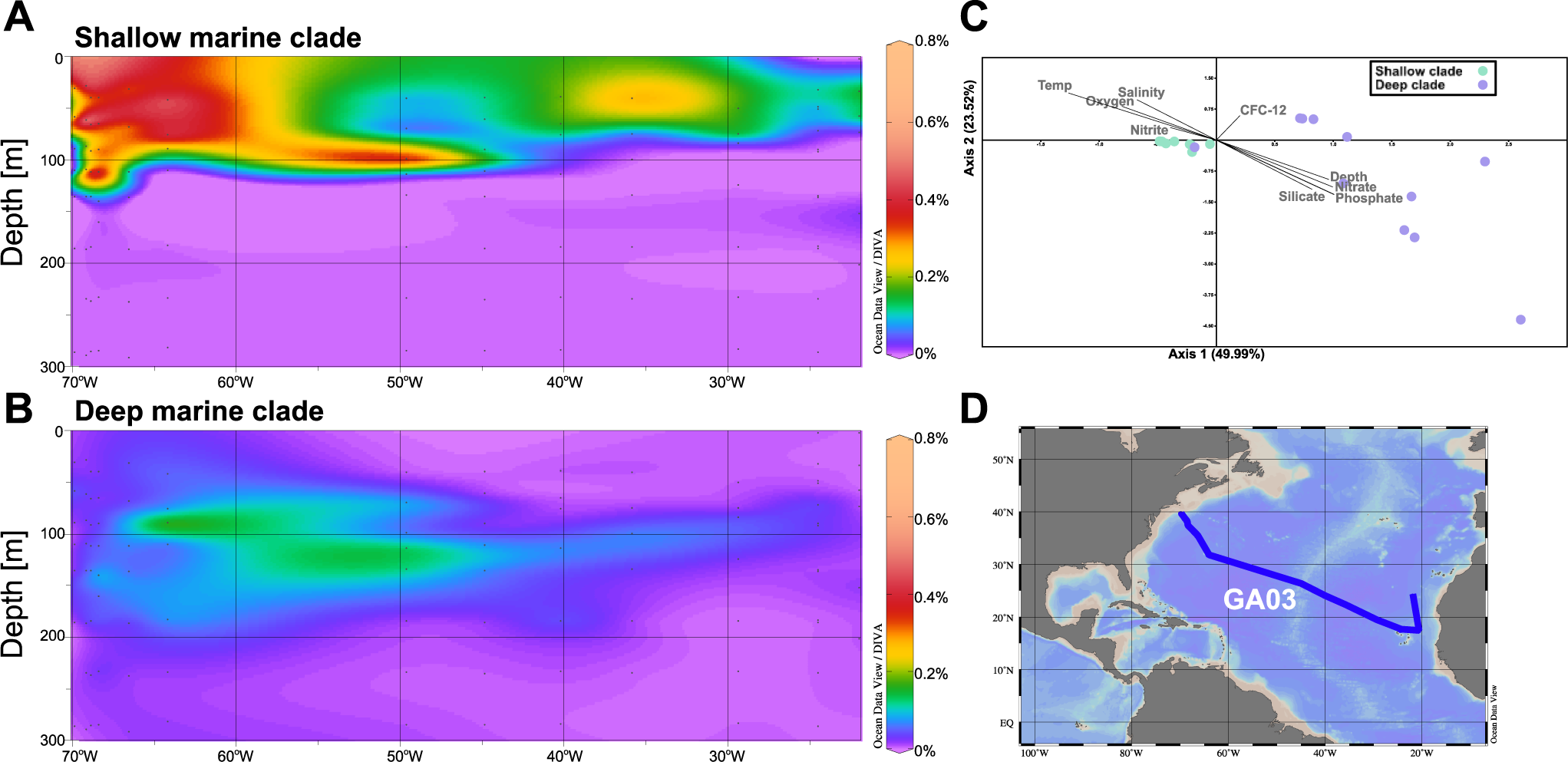
A. Ocean Data View plot of percent relative fraction for the *Dadabacteria* shallow clade along the GEOTRACES transect GA03. B. Ocean Data View plot of percent relative fraction for the *Dadabacteria* deep clade along the GEOTRACES transect GA03. C. Canonical correspondence analysis of the nonredundant marine *Dadabacteria* MAGs. Vectors denote correlations with environmental parameters and have been modified for easier visualization: trioplot amp 1.5, scaling type 2. D. Cruise track of GA03.

As exemplified by the GA03 cruise track in the North Atlantic, the resolution provided by bGT reveals that the shallow and deep clades are dominant above and below ∼100m depth (∼1% light level), respectively, and that this niche transition can be sharp, with the shallow clade dropping to a negligible component of the microbial community at this partitioning depth (Figure 3; Supplemental Table 11). This relationship can be observed for the other three bGT cruise tracks with some localized variation where the deep clade can be found at the surface and the shallow clade can be found at 250m, but for many stations there remains a sharp divided between the two clades at ∼50-100m depth (Supplemental Figure 2). Canonical correspondence analysis (CCA) of the GA03 environmental parameters support this niche transition as a majority of the deep clade correlated with depth and depth-associated parameters (nutrients, temperature, *etc*.; Figure 2C). Similar correlations between depth-associated parameters and the marine clades are observed for the other cruise tracks (Supplemental Figure 4). As has been shown previously, deep euphotic zone blue-light proteorhodopsins are adapted to low light incidence and capture a limited amount of light at 75m (Wang *et al*., 2003), the division between the shallow, proteorhodopsin-encoding clade and the deep clade reflect an evolutionary selective pressure of maintaining a light-responsive protein apparatus at depth and establish depth-specific niche boundaries between the two marine clades.

## Conclusion

The *Dadabacteria* phylum is an understudied clade with a limited number of genomic representatives. The broad analysis of the four major clades represented among publicly available genomes reveals a broad range of heterotrophic organisms, putatively involved in the recycling of microbially-derived POM, such as peptidoglycan and phospholipids. The terrestrial clades appear to be facultative anaerobes capable of using alternative electron acceptors, while the marine clades appear to be obligate aerobes. The marine clades have genomic features indicating extensive genome streamlining evolutionary pressures that mirror their ecological distribution in oligotrophic environments. Genome streamlining theory is an important hypothesis for explaining the prevalence of small genomes among cosmopolitan microorganisms and the *Dadabacteria* represent a clear example of the theory in action. The two distinct marine clades are differentiated in metabolic potential by the presence of light-associated adaptations, such as proteorhodopsin, terpenes, and carotenoids, supporting an argument that shallow clade possess a photoheterotrophic lifestyle. These adaptations are reflected in the ecological distribution of these clades with depth-partitioned niches of distinct shallow and deep clades. The *Dadabacteria* have multiple transitions that are of interest for understanding evolutionary pressures and adaptations in different environments, including: terrestrial to marine transitions; high to moderate/low temperature transitions; and adaptations from organic rich to organic poor environments. Further studies and the expansion of available genomes for this clade may provide specific insights as to how these transitions occur and manifest in microbial genomes.

## Data Availability

Several of the MAGs (TOBG-EAC99, TARA-RED-00009, TOBG-IN994, TOBG-MED731, TOBG-MED713, and TOBG-SP357) used in this study and underwent manual curation originated from the *Tara* Oceans dataset and were never submitted to NCBI to avoid duplication in GenBank. These curated MAGs are noted in Supplemental Table 1 and are available here: 10.6084/m9.figshare.12344207. As noted in Supplemental Table 1, MAGs with corresponding submissions in NCBI GenBank have been updated.

## Contributions

Analyses were conducted by E.D.G and B.J.T. Specifically, E.D.G. performed quality assessments, manual improvement of the MAGs, reconstructed the phylogeny, and recruitment procedure to determine ecological distributions. B.J.T. performed analyses related to functional annotations and genome streamlining. E.D.G and B.J.T. wrote the manuscript. The study was conceived by B.J.T.

## Supplemental Material

*(Prior to publication supplemental figures, tables, and data are available here: https://doi.org/10.6084/m9.figshare.12543488.v1)*

Supplemental Figure 1. Bubble plot of *Tara* Oceans sites and samples that recruit ≥0.05% relative fraction against the *Dadabacteria* MAGs. Bubble sizes scale from 0.051-0.471%.

Supplemental Figure 2. Bar plot of the number of *Tara* Oceans samples from the three depths and four filter fractions targeted as part of the expedition. Bars are filled in according to the number of samples that had ≥0.05% of the metagenome recruit to the MAGs of the shallow or deep *Dadabacteria* clades.

Supplemental Figure 3. Percent relative abundance of shallow and deep *Dadabacteria* clades displayed over the length of the three bioGEOTRACES cruise tracks (displayed in the corresponding maps).

Supplemental Figure 4. Canonical correspondence analysis of shallow or deep *Dadabacteria* MAGs for three individual bioGEOTRACES cruise tracks and all four cruise tracks combined. Vectors denote correlations with environmental parameters and have been modified for easier visualization: trioplot amp 1.5, scaling type 2.

Supplemental Table 1. Information for genomes used in this study, including source, phylogenetic assignment, genomes statistics, and values used in Figure 1.

Supplemental Table 2. Raw KEGG-Decoder output converting KEGG KO assignments to metabolic pathways of interested. Values from 0-1.

Supplemental Table 3. Raw counts for extracellular peptidases detected in *Dadabacteria* MAGs. Tab 1. All extracellular peptidases detected. Tab 2. A subset of extracellular peptidases displayed in Figure 1.

Supplemental Table 4. Raw counts for carbohydrate active enzymes detected in *Dadabacteria* MAGs. Tab 1. All carbohydrate active enzymes detected. Tab 2. A subset of carbohydrate active enzymes common in multiple MAGs.

Supplemental Table 5. eggNOG-mapper results for *Dadabacteria* MAGs. Tab 1. Terrestrial clade results. Tab 2. Hot Spring clade results. Tab 3. Shallow clade results. Tab 4. Deep clade results. Tab 5. Putative luciferases for all clades. Tab 6. Putative CAS proteins for all clades. Tab 7. Putative ABC-transporter components for all clades. Tab 8. eggNOG matches used in Figure 1.

Supplemental Table 6. A subset of functions of interest from KEGG and eggNOG determined for each of the four *Dadabacteria* clades.

Supplemental Table 7. Assignment of detected rhodopsins in the *Dadabacteria* shallow clade. Supplemental Table 8. AntiSMASH results for all *Dadabacteria* MAGs.

Supplemental Table 9. Percent relative fraction values from all marine metagenome used to assess the distribution of marine *Dadabacteria* MAGs.

Supplemental Table 10. RPKM values from all marine metagenome used to assess the distribution of marine *Dadabacteria* MAGs.

Supplemental Table 11. Input values used to generate ODV plots for Figure 2 and Supplemental Figure 2.

Supplemental Data 1. Newick file used to construct the phylogenetic tree in Figure 1 using the BAC120 markers that are part of GTDB-Tk.

Supplemental Data 2. Protein FASTA file of rhodopsins detected in *Dadabacteria* MAGs.

Supplemental Data 3. FASTA of the aligned region used to determine function and spectral tuning.

Supplemental Data 4. GEOTRACES combined and linked for metagenomes that are part of bioGEOTRACES.

## Notes

https://doi.org/10.6084/m9.figshare.12543488.v1

### Competing Interest Statement

The authors have declared no competing interest.

